# Detecting Signals of Seasonal Influenza Severity through Age Dynamics

**DOI:** 10.1101/016105

**Authors:** Elizabeth C. Lee, Cécile Viboud, Lone Simonsen, Farid Khan, Shweta Bansal

## Abstract

Background: Measures of population-level influenza severity are important for public health planning, but estimates are often based on case-fatality and case-hospitalization risks, which require multiple data sources, are prone to surveillance biases, and are typically unavailable in the early stages of an outbreak. In this study, we develop a severity index based on influenza age dynamics estimated from routine surveillance data that can be used in retrospective and early warning contexts.

Methods and Findings: Our method relies on the observation that age-specific attack rates vary between seasons, so that key features of the age distribution of cases may be used as a marker of severity early in an epidemic. We illustrate our method using weekly outpatient medical claims of influenza-like illness (ILI) in the United States from the 2001 to 2009 and develop a novel population-level influenza severity index based on the relative risk of ILI among working-age adults to that among school-aged children. We validate our ILI index against a benchmark that comprises traditional influenza severity indicators such as viral activity, hospitalizations and deaths using publicly available surveillance data. We find that severe influenza seasons have higher relative rates of ILI among adults than mild seasons. In reference to the benchmark, the ILI index is a robust indicator of severity during the period of peak epidemic growth (87.5% accuracy in retrospective classification), and may have predictive power during the period between Thanksgiving and the winter holidays (57.1% accuracy in early warning). We further apply our approach at the state-level to characterize regional severity patterns across seasons. We hypothesize that our index is a proxy for severity because working-age adults have both pre-existing immunity to influenza and a high number of contacts, infecting them preferentially in severe seasons associated with antigenic changes in circulating influenza viruses. Our analysis is limited by its application to seasonal influenza epidemics and a relatively short study period.

Conclusions: Our severity index and research on the link between age dynamics and seasonal influenza severity will enable decision makers to better target public health strategies in severe seasons and improve our knowledge of influenza epidemiology and population impact. These findings demonstrate that routine surveillance data can be translated into operational information for policymakers. Our study also highlights the need for further research on the putative age-related mechanisms of severity in seasonal and pandemic influenza seasons.

## 1 Introduction

The causes and characterization of population-level severity are crucial aspects to understanding influenza epidemiology and designing effective surveillance and control programs. Many mechanisms for variation in flu season severity are posited, but these findings have not been synthesized across fields. Environmental factors, like colder winters, are associated with severity, perhaps due to immune disregulation or the co-circulation and co-infection with other respiratory pathogens [1, 2]. Influenza virus genetic and antigenic data suggests that severity is linked to hemagglutinin protein novelty [3], and A/H3-dominant seasons have greater mortality rates than A/H1 or B dominant seasons [4, 5, 6, 7] as H3 strains disproportionately affect older populations [8]. Epidemiological studies of seasons following pandemics suggest that novel antigenic strains shift the expected age-related immune landscape, which generates relatively high susceptibility among adults [9, 10, 11, 12, 5, 13]. In congruence with a link between severity and low pre-existing immunity, high T cell concentrations correlate with subsiding illness and high viral shedding correlates with high clinical severity[14].

While causes of severity are not completely understood, current discourse about population-level seasonal influenza severity ties itself to experiences of severe patient-level outcomes. The Centers for Disease Control and Prevention (CDC) estimates a range of 3,000 to 49,000 influenza-associated all-cause deaths per year and over 200,000 hospitalizations in the United States during the period between 1976 and 2007 [7, 15], and generally characterizes seasonal severity through influenza-related hospitalization rates and mortality due to pneumonia and influenza, which provide a population-level view of patient-level outcomes (Figure 1). Clinical studies similarly focus on patient-level outcomes, where physicians use scoring techniques to rate overall patient severity or the severity of specific symptoms [16, 17].

**Figure 1:**
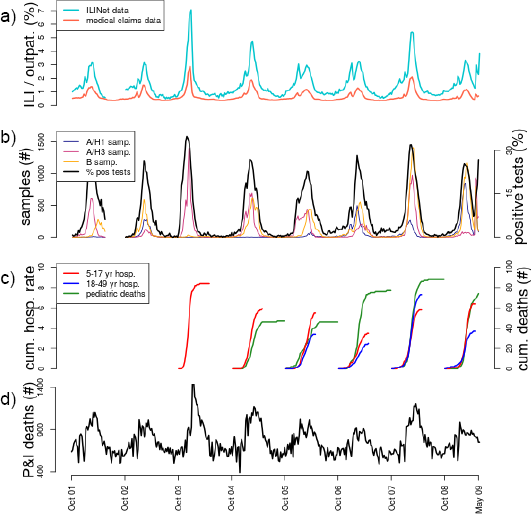
Influenza surveillance data over the 2001-02 to 2008-09 seasons. Characterization of ILI activity from October 2001 to May 2009 as a function of: a) ILI as a percentage of all outpatient visits in CDC’s ILINet and IMS Health medical claims data, b) influenza subtype samples and percentage of laboratory-confirmed influenza specimens, c) laboratory-confirmed influenza surveillance: cumulative hospitalization rates per 100,000 population for 5-17 year olds and 18-49 year olds, and cumulative pediatric deaths (under 18 years old) over the course of the season, and d) number of deaths attributed to pneumonia and influenza. All data except medical claims ILI are publicly available from CDC surveillance systems. Surveillance data on positive percentage of influenza tests, hospitalization rates, pediatric deaths, and pneumonia and influenza deaths are used to construct the benchmark.

Epidemiological analyses extrapolate proxies of patient-level severity such as case-fatality and case-hospitalization risk estimates to the population-level, but these metrics focus heavily on pandemic and emerging outbreaks [18, 19, 20, 21, 22]. Other studies model the relationship between excess mortality and morbidity rates [23, 24] or threshold excess P&I mortality rates in order to identify and detect severe flu seasons, but these metrics capture only one facet of the experience of flu across the population [4, 25]. Moreover, P&I mortality data is neither available in real-time nor collected by many national flu surveillance systems (e.g. in the European Union). In the past, the CDC used the Pandemic Severity Index, a pandemic rating system calibrated to case fatality ratios across the United States, to classify pandemic flu severity [26]. More recently, the CDC adopted a population-level severity framework that incorporates both clinical severity and transmissibility metrics, but the clinical severity component remains closely tied to case-fatality and similar ratios in the epidemiological data context [20].

In this work, we develop novel severity assessment metrics that synthesize existing information on viral activity, hospitalizations, and deaths, and explore proxies of population-level flu severity by examining age patterns influenza-like-illness (ILI) among the healthiest and largest segments of the population. Using a high coverage outpatient ILI dataset based on medical claims data from the United States, we introduce a retrospective index of flu severity, which is measured during the influenza epidemic growth period, to aid in epidemiological analysis and evaluation of public health responses. In addition, we show that this measure can be used for early warning, which can help decision makers tailor communication strategies regarding vaccination and antiviral usage, and enable corporate pharmacies to shift antiviral distribution to suit regional needs. Lastly, we carry out a spatial analysis of influenza severity to identify regions of the U.S. that represent or deviate from national-scale influenza severity to guide future influenza surveillance design.

## 2 Results

We examined weekly medical claims data of influenza-like illness (ILI) outpatient cases in the United States provided by IMS Health for the 2001-02 through 2008-09 flu seasons (Figure 2a). ILI reports were adjusted for increasing coverage in surveillance systems over time and age and region-specific estimates of health-care seeking behavior for ILI (See Methods). Recent analysis finds that ILI claims data accurately capture weekly fluctuations in influenza activity and season level intensity at high resolutions by age group and geographic location [27] and can be used to monitor the spatial spread of the disease [28].

**Figure 2:**
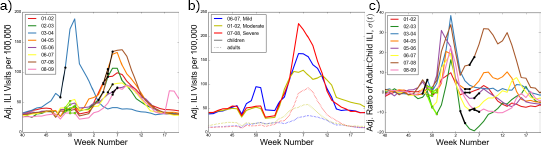
Influenza age dynamics follow different patterns than overall epidemic dynamics. (a) Medically attended outpatient ILI visits per 100,000 for the 2001-02 through 2008-09 flu seasons, adjusted for increasing surveillance data coverage and ILI care-seeking behavior, are displayed. The national early warning and retrospective classification periods are highlighted in green and black, respectively. (b) Child and adult ILI rates in mild, moderate and severe seasons (as identified by the benchmark index, *β*. (c) The normalized relative risk of adult ILI to child ILI rates (*σ*(*t*)), a proxy of age-specific disease burden, follows a regular seasonal pattern during the U.S. Thanksgiving and winter holiday periods, and diverges during the typical epidemic growth periods of January and February (around weeks 2-7).

In this study, we motivate and develop a metric for estimating seasonal influenza severity in the United States for an accurate, retrospective classification and an early warning context. To validate this metric, we also develop a benchmark metric based on traditional measures of severity derived from CDC surveillance sources. We then extended this approach to the state-level to discern regional severity patterns across our study period.

### 2.1 Exploring correlates of influenza severity

We developed a composite benchmark measure for influenza severity each season (*β*), which synthesizes CDC surveillance data on influenza-related hospitalizations and deaths from 1997-98 throughout 2013-14, excluding the 2009-10 season due to the H1N1 pandemic. The benchmark identified 2000-01, 2002-03, 2005-06, 2006-07, 2008-09, and 2011-12 as mild seasons, 1997-98, 1998-99, 2001-02, 2004-05, and 2007-08 as moderate seasons, and 1999-00, 2003-04, 2010-11, 2012-13, and 2013-14 as severe seasons. To identify possible drivers of seasonal severity, we examined Pearson’s correlation coefficients between *β* and covariates representing circulating subtypes, age-specific and total ILI rate from CDC’s ILINet, and temperature and precipitation (Supplement Figure 6). We find that adult and total ILI attack rates were positively correlated with *β*, and were the only correlations that achieve statistical significance at the alpha = 0.05 level.

Indeed, in examining the age-specific outpatient medical claims data, we found that peak ILI rates in children were always higher than that of working-age adults in a given season, and that they did not appear to correspond with rank order severity. On the other hand, adult peaks were higher in known severe seasons than milder ones (Figure 2b). This age-specific pattern was also observed when considering ILI rates in excess of a seasonal baseline (Supplement Figure 9).

The singling out of working-age adult ILI among severity correlates, prior work on the dynamic immune landscape in adults and children in shifting disease burdens with pandemic flu [9], and substantial interest in the literature in the roles of children and adults as the largest and most active segments of the population, motivate us to study influenza severity through the proxy of these two healthiest age groups.

### 2.2 Measuring severity through age-specific risks

We examined severity through a weekly proxy of age-specific disease burden, *σ*(*t*), which is a normalized relative risk of adult to child ILI rates at week *t*. (See Methods for details.) Severe seasons had relatively large adult disease burdens (high *σ*(*t*)) during the peak flu season, while mild seasons had relatively large child disease burdens (low *σ*(*t*)) during this time (Figure 2c). We used *σ*(*t*) to define measures of severity for the purposes of early warning in the period between Thanksgiving and Christmas 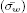, and retrospective classification during the primary epidemic growth period 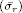, which were the only two windows with significant severity detection capabilities (Supplement Figure 4). We highlight that our metric does not measure transmissibility but is based on the age structure of cases.

We validated our relative risk severity measures with qualitative severity categorizations in the CDC flu season summaries, and quantitative classifications such as the benchmark (*β*) and other traditional severity metrics. Retrospective severity 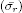 derived from the medical claims identified 2002-03, 2005-06, 2006-07, and 2008-09 as mild seasons, 2001-02 as a moderate season, and 2003-04, 2004-05, and 2007-08 as severe seasons (Pearson correlation coefficient of 0.77 and 87.5% accuracy when compared to *β*) (Figure 3a). The early warning index 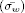 identified 2001-02, 2002-03 and 2006-07 as mild, 2004-05 and 2008-09 as moderate, and 2005-06 and 2007-08 as severe (Pearson correlation coefficient of 0.46 and 57.1% accuracy when compared to *β*) (Figure 3b). Among other severity metrics, hospitalization rates and total attack rates were positively correlated with 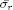 (Supplement Figure 12).

**Figure 3:**
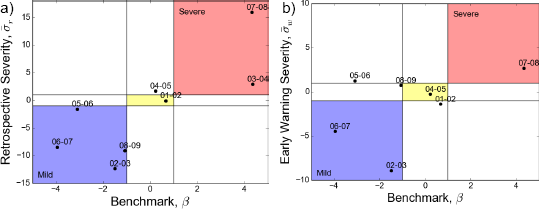
The retrospective severity index is an accurate measure of population-level severity, while the early warning index provides some predictive value. (a) Retrospective severity 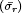 has a strong positive correlation (R = 0.77) and agrees with the benchmark for seven of eight seasons. (b) Early warning severity 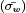 has a moderate positive correlation (R= 0.46) and agrees with the benchmark for four of seven seasons. The 2003-04 season was removed because it was an early flu season and the early warning period occurred after the retrospective period.

We also calculated our severity measures for CDC’s ILINet surveillance data and found that retrospective severity was well correlated between ILINet and the medical-claims-derived index (Pearson correlation coefficient of 0.775), but the ILINet index had a relative tendency to underestimate severity classifications (Supplement Figure 11). Also, ILINet 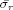 was well correlated with the benchmark (Pearson correlation coefficient of 0.70 and 62.5% accuracy), but 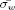 did not provide predictive value (Pearson correlation coefficient of -0.26 and 15.4% accuracy) (Supplement Figure 10).

### 2.3 Spatial patterns in influenza severity

We examined state-level patterns in severity by calculating retrospective and early warning indexes from age-specific ILI rates at this higher resolution spatial scale (details in the Supplementary Methods). Regardless of classification at the nation-level, severity ranged from mild to severe at the state-level in a single season (Figure 4a). Across the eight study seasons, the mid-Atlantic states of Virginia and North Carolina experienced significantly more severe seasons than the national 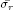 (interquartile range of deviation between national and state-level 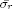 is above zero) (Figure 4b). No state had consistently milder flu seasons, but evidence suggests that the western United States may have milder seasons than the national 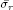.

**Figure 4:**
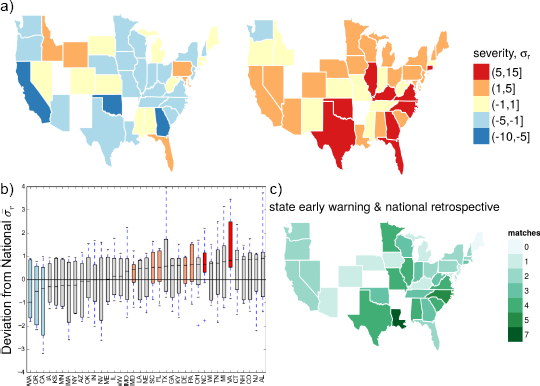
State-level patterns of seasonal flu severity. a) State retrospective severity classifications may range from mild to severe in a single season regardless of the national severity classification (left panel, 2006-07 mild season; right panel, 2007-08 severe season). (b) Deviation between state-level and national 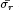 across the eight study seasons, where states with greater severity have positive values and vice versa. North Carolina and Virginia have significantly more severe seasons (interquartile range above zero), and other Eastern states like Delaware, Pennsylvania, Maryland, South Carolina, and Florida regularly have more severe seasons than the rest of the U.S.. Western states – Washington, California and Oregon – have more milder seasons on average. Only states with retrospective severity indexes for all eight study seasons are reported in b. (c)Number of matches between a given state’s early warning severity classification and the national retrospective classification. Louisiana matched its early warning signal with national retrospective severity for seven of eight seasons, but this result did not deviate significantly from expectation had severity been randomly assigned. Only the 36 states with retrospective and early warning severity indexes for all seasons are reported in c.

We examined whether there were “sentinel” states, which may correctly indicate national or state-level retrospective severity during the early warning period. In Figure 4c, we counted matching state-level 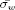 and nation-level 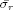 severity among the 36 states with classifications available for the entire study period. Louisiana’s early warning severity matched the national retrospective severity for seven of eight seasons, suggesting that it may serve as a reasonable sentinel, but this result was not statistically different than expectation had severity been randomly assigned to states. We also examined state-level sentinels (Supplement Figure 13c), but found no states with significant predictive capability (details in the Supplementary Methods).

## 3 Discussion

Our severity assessment index detects indirect signals of seasonal influenza severity from a single-source, real-time ILI dataset with obligatory reporting, and it is flexible enough for use in retrospective and early warning contexts and with other ILI surveillance systems such as CDC’s ILINet. Motivated by our finding that adult ILI visits were highly correlated with traditional measures of severity (including hospitalization and deaths), we developed a proxy for influenza severity based on the ratio of ILI risk among adults relative to that among children. As school-aged children and adults are the lowest risk groups for seasonal influenza complications and death [29], our metric seeks to measure signals of severity indirectly through populations that are well-represented in influenza case data and well-connected to high-risk populations. To validate this metric, we constructed a novel benchmark of severity from public data that comprises traditional severity indicators. With respect to this benchmark, our severity index performs with 88% accuracy retrospectively, and with 57% accuracy prospectively, with mis-classifications that err conservatively from the standpoint of public health (i.e., they tend to be false positives instead of false negatives). When we extended our severity analysis to the state-level, we found evidence for more severe influenza seasons than the national average in the mid-Atlantic region of the U.S., and for the possibility that certain states (Louisiana was identified in our analysis) act as “sentinels” for severity in the national epidemic.

The retrospective classification of influenza severity can inform public health systems evaluations and enables historical analysis of severe season attributes, which will improve our understanding of influenza disease ecology. Early warning of severity (9-12 weeks before typical epidemic peak) enables public health agencies to tailor communication campaigns for vaccination and antiviral usage. These health communication campaigns from official information sources regarding pharmaceutical [30, 31] and behavioral [32, 33] interventions for influenza have been shown to improve individual health-related behaviors, and could have an impact on epidemic outcomes. Finally, our metrics are flexible enough to be used with different geographic scales and ILI surveillance systems, and their broad application may enable the multi-scale coordination of influenza public health policy. In Figure 5, we map our severity metric to functional indicators of influenza burden, including peak ILI, hospitalization, and mortality rate to provide translation of our metric in an operational context.

**Figure 5:**
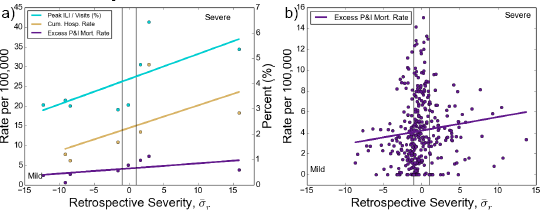
Translation of the retrospective severity index 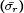 to operational public health indicators of influenza’s burden on the health care system. a) At the national scale, a severe 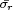 categorization translates approximately as 30-45% of peak week outpatient visits due to ILI (ILINet), 10-20 confirmed influenza-related hospitalizations per 100,000 over the course of the season, and seasonal excess P&I mortality rates of 3-10 per 100,000 (Pearson’s R = 0.68, 0.59, and 0.54 respectively). b) At the state level, a severe 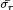 categorization corresponds to approximately 0 to 10 excess P&I mortality rates (Pearson’s R = 0.16). It appears that moderate 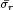 seasons have an inclusively larger range of excess P&I mortality rates, which suggests that moderate seasons at the national level represent a mixture of severe and mild flu seasons at the state-level.

While the elderly are a traditional high-risk group due to their high mortality and the prevalence of comorbidities that increase the likelihood of flu-related complications [6], this age group is less likely to experience illness overall due to increased prior immunity and weaker contact patterns [34]. Instead, our index looks for indirect signals of severity using the age dynamics of “healthier” populations, reasoning that seasons with larger effects in children and adults are more severe for the entire population. We posit that school-aged children experience substantial flu morbidity every season because they have high numbers of potential disease-causing contacts [35, 36, 37] and greater susceptibility due to less prior exposure to influenza and less developed immune systems. Adults have fewer contacts and greater prior immunity than children, so they experience high flu activity only when the flu season is severe, regardless if the cause is strain novelty, higher transmissibility, greater virulence, or some combination of factors. High connectivity between adults and other age groups [36] and the role of adults in seeding new regions [38, 39] may explain why seasons that burden adult populations are also severe for the entire population. In addition, A/H3 seasons, which cause more severe seasons due to immune escape [40, 41], are thought to affect adults more than children [8, 42]. (Our metric, however, detects severity for both H1 and H3-dominant seasons (Supplement Figure 14)). Notably, our two severity classification periods occur during the first half of the epidemic, suggesting that time-varying contact patterns rather than prior immunity, which presumably affects the entire epidemic, better explains the severity signal detection witnessed by our index (Supplement Figure 4). The retrospective severity index captures age dynamics during the peak flu season, while the combination of changes to contact patterns and reductions in care-seeking behavior during the holidays allow the early warning index to detect signals during a “severity testbed” period in the United States. Future analyses should disambiguate the age patterns characterized in our study as a harbinger or result of population-level severity and examine the hypothesis that holiday contacts seed broader infection in different age groups [43] or new locations.

We note that the early warning severity index is limited in its accuracy. In some seasons, circulation of flu is low around the Thanksgiving and winter holiday periods, and other respiratory pathogens like respiratory syncytial virus (RSV), rhinoviruses, and Haemophilus influenzae may confound the adult and child influenza-like illness dynamics used in our metric [44]. On the other hand, flu seasons that arrive earlier than usual and peak during this holiday period (eg. 1999-2000, 2003-04, 2013-14) also pose a problem for our early warning signal because there is a strong shift in age dynamics every season towards adult disease burden during the winter, and to a lesser extent, Thanksgiving holidays. Nevertheless, the retrospective severity signal identifies those early seasons as severe, thus corroborating findings that early seasons are associated with increased severity, perhaps due to increased population susceptibility to antigenically-novel strains [45]. The early warning index for ILINet surveillance did not perform well as compared to the medical claims data, but this may be explained by ILINet’s smaller sample size and narrower syndromic definition of flu, both of which could limit the detection of influenza activity during the early warning period (Supplement Figure 10). Lastly, the early warning index may be limited to use in the United States; shifting contact patterns and holiday-related care-seeking behavior during the Thanksgiving holiday may improve early warning signal detection (Supplement Figure 16).

The application of our severity index to the state-level builds upon our current knowledge of spatial influenza epidemiology. The western U.S. had the mildest absolute flu activity based on the 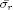 metric, corroborating other studies (Supplement Figure 13a) [46], but additional independent validation of state-level severity results is needed. Both the state-level severity index and state-level excess P&I mortality rates illustrate large within-season variation and states should expect the greatest heterogeneity in influenza-related mortality during moderate seasons (Figures 4 and 5b). Future studies may reveal that this spatial variation in severity can be explained by different regional subtype circulation, pre-existing immunity, age distributions, or vaccine coverage rates (Supplement Figure 15).

Further research is needed to improve the application of our severity metric to pandemic influenza. Young children and the elderly contribute most to mortality in seasonal flu, while pandemic scenarios are notable for their tendency to generate greater mortality or serious complications among groups that do not normally experience strong illness, such as adults under 65 [5, 18, 47, 48, 42]. These events are characterized by different distributions of age risk: an initial pandemic wave may be dominated by morbidity among school-aged children, and some empirical and modeling studies suggest that adults are more likely to become infected in the season following a pandemic [9, 10, 11, 12, 13]. While the mechanisms for this remain poor understood, it has been suggested that there is an accumulation of heterologous immunity with age [49]. Additionally, it has been observed that excess P&I mortality shifts from the elderly to younger ages in the years following a pandemic, perhaps signaling shifted dynamics between children and adults during this time as well. Our index would not capture severity in the first and second waves of pandemic virus circulation, which is why we exclude the 2009-10 season from our analysis, and unstable age dynamics in post-pandemic seasons may explain misclassifications of recent seasons from ILINet data (Supplement Figure 10 c&d).

Our severity index relies on real-time age-specific incidence data, which has not been utilized thus far in other “big data”-based surveillance (eg. Google Flu Trends), apart from questionable relationships between clinical data and flu-related big data [50]. Traditional ILI surveillance in the form of CDC’s ILINet does provide real-time age-specific data, but the use of electronic medical claims data has advantages. ILINet comprises voluntary reports from participating physicians, so competing time demands may skew ILI reports. Medical claims data is a more obligatory form of health care provider reporting, captures ILI activity at least as well as traditional surveillance, may be examined in real-time, and provides higher coverage, greater spatial resolution, and finer age-specific disease information due to its insurance record nature [27]. Medical claims and ILINet data are both subject to physician biases regarding the demographics and seasonality of influenza and doctor’s office closures. They also have healthcare-seeking behavior biases; school-aged children have higher rates of healthcare-seeking behavior for ILI than adults (approximately 1.1 to 1.4 times higher) [51, 52, 53], which is why we adjust for these biases in both datasets (See Supplementary Methods). Nevertheless, differences in surveillance data biases and study period may explain scaling issues with retrospective classification and poor performance in early warning severity for ILInet data (Supplement Figure 10 c&d) Additional studies on age-specific disparities in health care insurance and access to care, especially in consideration of ongoing changes to the U.S. health care system, are needed to better quantify biases in medical claims data and our index.

The correlation of the retrospective index with an independent, composite severity benchmark and a multitude of other severity indicators, in addition to its consistent interpretation across seasons, suggests a relationship with population-level seasonal flu severity. As far as we know, this study is the first to link population-level severity with age dynamics in a broad sense, thus representing proof of concept that influenza age-related disease dynamics may provide epidemiological understanding beyond the information provided at face value.

## 4 Materials and Methods

### 4.1 Ethics statement

Patient records and information in the medical claims dataset were anonymized, de-identified, and aggregated by IMS Health, which handles data that is routinely collected for health insurance purposes. Mr. Farid Khan, Director of Advanced Analytics at IMS Health, granted the researchers access to the data. While the database is not accessible online, interested researchers should refer to the IMS Health website: http://www.imshealth.com/portal/site/imshealth. All analyses were performed with aggregated time series data for influenza-like illness rather than patient-level information, and similar to other epidemiological analyses of administrative insurance data, no institutional review board approval was sought.

### 4.2 ILI surveillance data

Our analyses utilized weekly physician’s office and outpatient visit data from October 2001 to May 2009 for influenza-like illness from a records-level database of CMS-1500 medical claims. Total visits captured by this commercially-available medical claims dataset increased by approximately 3.5 times over the duration of our study as more data streams were purchased over time; by 2009, the data comprises 934 three-digit zipcode prefixes and documents 62% of all visits to physicians (IMS health collected data from 354,402 of 560,433 active physician practices in the US) in the United States [28, 27]. A previous study derived a synthetic definition of ILI based on International Classification of Diseases (ICD-9) codes – influenza (487-488) or [fever and (respiratory symptoms or febrile viral illness) (780.6 and (462 or 786.2))] or prescription of oseltamivir (most commonly, 079.99) – and used this definition to validate the medical claims data to patterns from CDC’s ILINet (details in Supplementary Methods) [27]. We consider the population of school-age children as 5-19 years old and working-age adults as 20-59 years old. Seasonal flu epidemics occur at varying times during the *flu season* enduring from October through May, which is roughly week 40 in one year to week 20 in the subsequent year. To test the suitability of our index for a different ILI surveillance system and a larger range of seasons, we use CDC’s publicly available ILINet data, which is available from October 1997 to May 2014; by 2014, the data documents outpatient visits from approximately 1,900 health care providers in the U.S. each week. We adjust the total volume of ILI cases to account for increasing coverage of the surveillance systems each year and for different spatial and age-specific estimates of health-care seeking behavior for ILI for both medical claims and ILINet surveillance data (See Supplementary Methods). All reported ILI rates are adjusted for increasing system coverage in their respective datasets and the reported estimates of health-care seeking behavior by age for ILI [51, 54].

### 4.3 Defining a severity benchmark

To create a synthetic composite severity *benchmark* (*β*), we aggregate CDC surveillance data on positive laboratory confirmations of influenza infections, P&I deaths, laboratory-confirmed influenza deaths in children under 18 years, and hospitalization in five to seventeen year olds and eighteen to forty-nine year olds to the flu season level and z-normalize each metric across the seasons (Figure 1). The z-normalized value for each criterion is summed to generate *β*. Surveillance systems that are not available for certain seasons in the specified time frame do not contribute to the index during that season (Supplement Table 2). *β* < –1 is classified as a mild season according to the benchmark, *β* > 1 as a severe season, and –1 ≤ *β* ≤ 1 as moderate. All of these factors appear to have a strong positive relationship with the synthetic benchmark *β*, indicating that no single, unique metric corresponds with severity (Supplement Figure 1). Sensitivity analyses for different normalization periods and data components for the benchmarks of the medical claims and ILINet data are performed in the Supplementary Methods and Supplement Figure 2. It should be noted that *β* classifications differ between the 2001-2009 study period and the 1997-2014 data that are used in some analyses in this paper because the values are normalized across all years. Consequently, this benchmark is relative to the severity of other seasons from which it is composed.

Since the 2009 H1N1 pandemic, testing practices in the United States have changed, including the more frequent use of rapid influenza-diagnostic tests and reverse-transcription polymerase chain reaction (RT-PCR) tests and the increased practice of pre-screening respiratory specimens by public health laboratories [55]. Consequently, the percent of positive lab confirmations may not serve as a valid indicator of seasonal influenza activity in a benchmark severity index that includes influenza seasons that occurring after 2009.

### 4.4 Exploratory correlation analysis

We examine pairwise Pearson cross-correlation coefficients of the benchmark severity index (*β*) over the 1997-98 to 2013-14 flu seasons (excluding 2009-10 due to the 2009 H1N1 pandemic) with nation-level covariates on viral strain distribution (the proportion of all positive influenza laboratory samples characterized as H3 at the end of the season) from World Health Organization (WHO) and National Respiratory and Enteric Virus Surveillance System (NREVSS) laboratories, age-specific and total disease burden (seasonal outpatient ILI rates of toddlers, children, adults, the elderly, and the total population) from CDC ILINet, and environmental variables (average temperature and precipitation during flu season months for 48 contiguous US states) from NOAA’s National Data Climate Center (NCDC). Pairwise tests of association between *β* and covariates of interest were deemed significant if they achieved *α* ≤ 0.05.

### 4.5 Retrospective and early warning severity

To construct our severity index, we first calculate the relative risk of (adjusted) ILI among adults that in school-aged children: 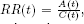, where *A(t)* and *C(t)* are the number of ILI cases captured in the surveillance system in a given week and spatial region (national or state) divided by the group’s population size in adults and school-age children, respectively. Since ILI is not restricted to laboratory-confirmed flu cases and baseline ILI activity varies from year to year, we transform the weekly RR time series through z-normalization to define a time series of severity as 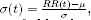 where *μ* is the mean and *σ* is the standard deviation of the *RR(t)* values during a specified baseline period. We define the *baseline period* as the first seven weeks of the flu season, the beginning of October to mid-November (weeks 40-46). This baseline overlaps with neither the typical peak flu season in the United States (January to March), nor the two classification periods in our severity framework, retrospective and early warning. We assess other baseline periods and find that the retrospective severity index is somewhat sensitive to summer baseline periods (Supplement Figure 3), but that our chosen period best represents baseline age dynamics (Supplement Figure 8).

Two severity classification periods are identified under our framework, the *retrospective* and the *early warning* periods. For most seasons, the retrospective period is the two week period that begins three weeks before the ILI peak in a given flu season ILI curve. The retrospective period is identified for the entire U.S. and each state (eg. In a given season, California’s retrospective severity is tied to peak ILI week in California and national retrospective severity precedes the peak ILI week at the national level). The early warning period is the two week period that begins two weeks after the Thanksgiving holiday in the United States. We define *retrospective severity* 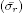 as the mean of *σ*(*t*) for weeks in the retrospective period and *early warning severity* 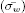 as the mean of *σ*(*t*) for weeks in the early warning period. The early warning and retrospective period 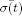 values are the only weeks over the year that correlate significantly with *β*, further supporting the uniqueness of these periods (Supplement Figure 4).

Retrospective severity provides a more accurate classification of seasonal severity because it captures the disease dynamics of the primary epidemic growth period, while the early warning severity provides an earlier assessment of severity between the Thanksgiving and winter holidays, which may serve as spatial seeding events. Seasons are classified mild if 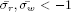, moderate if 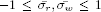 and severe if 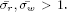 Severity may also be accurately estimated with the relative risk of ILI visits between adults and children over the entire epidemic period, but use of the two-week retrospective period is preferred due to scaling differences (Supplement Figure 5). Early warning severity is not calculated for early flu seasons (eg. 2003-04) and further detail may be found in the Supplementary Methods.

### 4.6 State-specific analyses

Deviation from the national baseline is calculated as the difference between state-level and national retrospective index values 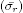, divided by the absolute value of national 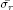 (Figure 4b). Sentinel state matches are shown only for states where retrospective and early warning classifications are available for the entire study period (Figure 4c).

### 4.7 Operational public health indicators

Percent of peak week outpatient visits due to ILI was identified using CDC ILINet data, and peak week is identified as the week in which the maximum of total ILI cases is achieved during a given season (Figure 5a).

We calculated state-level excess P&I mortality rates by state and season for the 2001-02 to 2008-09 study period from monthly pneumonia and influenza deaths at all places of death provided by the CDC WONDER Multiple Cause of Death database, which is publicly available online. As in Viboud et al. 2005, non-flu P&I mortality, which excludes the flu season period from from December to April, were fit with a Serfling-type regression model to identify the seasonal baseline. December to April P&I mortality in excess of the baseline were categorized as excess P&I mortality and aggregated to the season level [56]. National excess P&I mortality rates by season represent the population-weighted average of all state-level excess P&I mortality rates in that season.

## Acknowledgments

The authors thank Matthew Biggerstaff for providing data on age-specific ILI care-seeking behavior and valuable comments on our draft and Jason Asher for useful suggestions for the analysis. We also thank Réseau Sentinelles, Institut National de la Santé et la Recherche Médicale (INSERM), Université Pierre et Marie Curie (UPMC) [http://www.sentiweb.fr] for sharing age-specific influenza surveillance data for France as used for supplementary analyses. This work was supported by the RAPIDD Program of the Science & Technology Directorate, Department of Homeland Security and the Fogarty International Center, National Institutes of Health.

## References

[1] Hardelid P, Andrews N, Pebody R. Excess mortality monitoring in England and Wales during the influenza A(H1N1) 2009 pandemic. Epidemiol Infect. 2011 Sep;139(9):1431–9.

[2] Group TE. Cold exposure and winter mortality from ischaemic heart disease, cerebrovascular disease, respiratory disease, and all causes in warm and cold regions of Europe. Lancet. 1997;349:1341–1346.

[3] Wolf YI, Nikolskaya A, Cherry JL, Viboud C, Koonin E, Lipman DJ. Projection of seasonal influenza severity from sequence and serological data. PLoS Curr. 2010 Jan;2:RRN1200.

[4] Simonsen L, Clarke MJ, Williamson GD, Stroup DF, Arden NH, Schonberger LB. The impact of influenza epidemics on mortality: introducing a severity index. Am J Public Health. 1997 Dec;87(12):1944–50.

[5] Simonsen L, Clarke MJ, Schonberger LB, Arden NH, Cox NJ, Fukuda K. Pandemic versus epidemic influenza mortality: a pattern of changing age distribution. J Infect Dis. 1998 Jul;178(1):53–60.

[6] Thompson WW, Shay DK, Weintraub E, Brammer L, Cox N, Anderson LJ. Mortality Associated with Influenza and Respiratory Syncytial Virus in the United States. J Am Med Assoc. 2003;289(2):179–186.

[7] Thompson WW, Comanor L, Shay DK. Epidemiology of Seasonal Influenza: Use of Surveillance Data and Statistical Models to Estimate the Burden of Disease. J Infect Dis. 2006;194(Suppl 2).

[8] Khiabanian H, Farrell GM, St George K, Rabadan R. Differences in patient age distribution between influenza A subtypes. PLoS One. 2009 Jan;4(8):e6832.

[9] Bansal S, Pourbohloul B, Hupert N, Grenfell B, Meyers LA. The shifting demographic landscape of pandemic influenza. PLoS One. 2010 Jan;5(2):e9360.

[10] Dávila J, Chowell G, Borja-aburto VH, Viboud C, Muñiz CG. Substantial Morbidity and Mortality Associated with Pandemic A / H1N1 Influenza in Mexico, Winter 2013-2014: Gradual Age Shift and Severity. PLoS Curr Outbreaks. 2014;2014(October 2013).

[11] Gómez-Gómez A, Magaña Aquino M, Bernal-Silva S, Araujo-Meléndez J, Comas-García A, Alonso-Zúñiga E, et al. Risk Factors for Severe Influenza A, Related Pneumonia in Adult Cohort, Mexico, 2013 to 14. Emerg Infect Dis. 2014;20(9).

[13] Rahamat-Langendoen JC, Tutuhatunewa ED, Schölvinck EH, Hak E, Koopmans M, Niesters HGM, et al. Influenza in the immediate post-pandemic era: a comparison with seasonal and pandemic influenza in hospitalized patients. J Clin Virol. 2012 Jun;54(2):135–40.

[13] Skowronski DM, Hottes TS, Janjua NZ, Purych D, Sabaiduc S, Chan T, et al. Prevalence of seroprotection against the pandemic (H1N1) virus after the 2009 pandemic. CMAJ. 2010 Nov;182(17):1851–6.

[14] Wilkinson TM, Li CKF, Chui CSC, Huang AKY, Perkins M, Liebner JC, et al. Preexisting influenza-specific CD4+ T cells correlate with disease protection against influenza challenge in humans. Nat Med. 2012 Feb;18(2):274–280.

[15] Thompson MG, Shay DK, Zhou H, Bridges CB, Cheng PY, Burns E, et al. Estimates of Deaths Associated with Seasonal Influenza - United States, 1976 - 2007. Morb Mortal Wkly Rep. 2010;59(33).

[16] Fleming DM, Moult AB, Keene O. Indicators and significance of severity in influenza patients. Int Congr Ser. 2001;1219:637–643.

[17] Frank AL, Taber LH, Wells JM. Comparison of Infection Rates and Severity of Illness for Influenza A Subtypes H1N1 and H3N2. J Infect Dis. 1985;151(1):73–80.

[18] Presanis AM, De Angelis D, Hagy A, Reed C, Riley S, Cooper BS, et al. The severity of pandemic H1N1 influenza in the United States, from April to July 2009: a Bayesian analysis. PLoS Med. 2009 Dec;6(12):e1000207.

[19] Lipsitch M, Finelli L, Heffernan RT, Leung GM, Redd SC. Improving the evidence base for decision making during a pandemic: the example of 2009 influenza A/H1N1. Biosecurity Bioterrorism Biodefense Strateg Pract Sci. 2011 Jun;9(2):89–115.

[20] Reed C, Biggerstaff M, Finelli L, Koonin LM, Beauvais D, Uzicanin A, et al. Novel Framework for Assessing Epidemiologic Effects of Influenza Epidemics and Pandemics. Emerg Infect Dis. 2013;19(1):85–91.

[21] Garske T, Legrand J, Donnelly Ca, Ward H, Cauchemez S, Fraser C, et al. Assessing the severity of the novel influenza A/H1N1 pandemic. BMJ. 2009 Jan;339:b2840.

[22] Yu H, Cowling BJ, Feng L, Lau EHY, Liao Q, Tsang TK, et al. Human infection with avian influenza A H7N9 virus: an assessment of clinical severity. Lancet. 2013 Jul;382(9887):138–45.

[23] Denoeud L, Turbelin C, Ansart S, Valleron AJ, Flahault A, Carrat F. Predicting pneumonia and influenza mortality from morbidity data. PLoS One. 2007 Jan;2(5):e464.

[24] van den Wijngaard CC, van Asten L, Meijer A, van Pelt W, Nagelkerke NJD, Donker Ga, et al. Detection of excess influenza severity: associating respiratory hospitalization and mortality data with reports of influenza-like illness by primary care physicians. Am J Public Health. 2010 Nov;100(11):2248–54.

[25] Simonsen L, Clarke MJ, Stroup DF, Williamson GD, Arden NH, Cox NJ. A Method for Timely Assessment of Influenza-Associated Mortality in the United States. Epidemiology. 1997;8(4):390–395.

[26] HHS, CDC. Interim Pre-pandemic Planning Guidance: Community Strategy for Pandemic Influenza Mitigation in the United States. 2007;.

[27] Viboud C, Charu V, Olson D, Ballesteros S, Gog J, Khan F, et al. Demonstrating the Use of High-Volume Electronic Medical Claims Data to Monitor Local and Regional Influenza Activity in the US. PLoS One. 2014 Jan;9(7):e102429.

[28] Gog JR, Ballesteros S, Viboud C, Simonsen L, Bjornstad ON, Shaman J, et al. Spatial Transmission of 2009 Pandemic Influenza in the US. PLoS Comput Biol. 2014 Jun;10(6):e1003635.

[29] Sebastian R, Skowronski DM, Chong M, Dhaliwal J, Brownstein JS. Age-related trends in the timeliness and prediction of medical visits, hospitalizations and deaths due to pneumonia and influenza, British Columbia, Canada, 1998-2004. Vaccine. 2008 Mar;26(10):1397–403.

[30] Flood EM, Rousculp MD, Ryan KJ, Beusterien KM, Divino VM, Toback SL, et al. Parents’ decision-making regarding vaccinating their children against influenza: A web-based survey. Clin Ther. 2010 Aug;32(8):1448–67.

[31] Brewer NT, Chapman GB, Gibbons FX, Gerrard M, McCaul KD, Weinstein ND. Meta-analysis of the relationship between risk perception and health behavior: the example of vaccination. Heal Psychol. 2007 Mar;26(2):136–45.

[32] Park JH, Cheong HK, Son DY, Kim SU, Ha CM. Perceptions and behaviors related to hand hygiene for the prevention of H1N1 influenza transmission among Korean university students during the peak pandemic period. BMC Infect Dis. 2010 Jan;10:222.

[33] Timpka T, Spreco A, Gursky E, Eriksson O, Dahlström O, Strömgren M, et al. Intentions to perform non-pharmaceutical protective behaviors during influenza outbreaks in Sweden: a cross-sectional study following a mass vaccination campaign. PLoS One. 2014 Jan;9(3):e91060.

[34] Gefenaite G, Tacken M, Kolthof J, Mulder B, Korevaar JC, Stirbu-Wagner I, et al. Predictors of influenza in the adult population during seasonal and A(H1N1)pdm09 pandemic influenza periods. Epidemiol Infect. 2014 May;142(5):950–4.

[35] Kucharski AJ, Kwok KO, Wei VWI, Cowling BJ, Read JM, Lessler J, et al. The contribution of social behaviour to the transmission of influenza A in a human population. PLoS Pathog. 2014;.

[36] Mossong J, Hens N, Jit M, Beutels P, Auranen K, Mikolajczyk R, et al. Social contacts and mixing patterns relevant to the spread of infectious diseases. PLoS Med. 2008 Mar;5(3):e74.

[37] Wallinga J, Teunis P, Kretzschmar M. Using data on social contacts to estimate age-specific transmission parameters for respiratory-spread infectious agents. Am J Epidemiol. 2006 Nov;164(10):936–44.

[38] Viboud C, Bjornstad ON, Smith DL, Simonsen L, Miller MA, Grenfell BT. Synchrony, waves, and spatial hierarchies in the spread of influenza. Science (80-). 2006;312(5772):447–451.

[39] Apolloni A, Poletto C, Colizza V. Age-specific contacts and travel patterns in the spatial spread of 2009 H1N1 influenza pandemic. BMC Infect Dis. 2013 Jan;13:176.

[40] Ferguson NM, Galvani AP, Bush RM. Ecological and immunological determinants of influenza evolution. Nature. 2003;422(March):428–433.

[41] Chowell G, Miller Ma, Viboud C. Seasonal influenza in the United States, France, and Australia: transmission and prospects for control. Epidemiol Infect. 2008 Jun;136(6):852–64.

[42] Lemaitre M, Carrat F. Comparative age distribution of influenza morbidity and mortality during seasonal influenza epidemics and the 2009 H1N1 pandemic. BMC Infect Dis. 2010 Jan;10(April 2009):162.

[43] Eames KTD, Tilston NL, Edmunds WJ. The impact of school holidays on the social mixing patterns of school children. Epidemics. 2011 Jun;3(2):103–8.

[44] Falsey AR, Hennessey PA, Formica MA, Cox C, Walsh EE. Respiratory Syncytial Virus Infection in Elderly and High-Risk Adults. N Engl J Med. 2005;352(17):1749–1759.

[45] Boni MF, Gog JR, Andreasen V, Feldman MW. Epidemic dynamics and antigenic evolution in a single season of influenza A. Proc Biol Sci. 2006 Jul;273:1307–16.

[46] Wenger JB, Naumova EN. Seasonal synchronization of influenza in the United States older adult population. PLoS One. 2010 Jan;5(4):e10187.

[47] Louie JK, Jean C, Acosta M, Samuel MC, Matyas BT, Schechter R. A review of adult mortality due to 2009 pandemic (H1N1) influenza A in California. PLoS One. 2011 Jan;6(4):e18221.

[48] Lemaitre M, Carrat F, Rey G, Miller M, Simonsen L, Viboud C. Mortality burden of the 2009 A/H1N1 influenza pandemic in France: comparison to seasonal influenza and the A/H3N2 pandemic. PLoS One. 2012 Jan;7(9):e45051.

[49] Epstein SL. Prior H1N1 influenza infection and susceptibility of Cleveland Family Study participants during the H2N2 pandemic of 1957: an experiment of nature. J Infect Dis. 2006 Jan;193(1):49–53.

[50] Olson DR, Konty KJ, Paladini M, Viboud C, Simonsen L. Reassessing Google Flu Trends data for detection of seasonal and pandemic influenza: a comparative epidemiological study at three geographic scales. PLoS Comput Biol. 2013 Jan;9(10):e1003256.

[51] Biggerstaff M, Jhung M, Kamimoto L, Balluz L, Finelli L. Self-reported influenza-like illness and receipt of influenza antiviral drugs during the 2009 pandemic, United States, 2009-2010. Am J Public Health. 2012;102(10):e21–26.

[52] Brooks-Pollock E, Tilston N, Edmunds WJ, Eames KTD. Using an online survey of healthcare-seeking behaviour to estimate the magnitude and severity of the 2009 H1N1v influenza epidemic in England. BMC Infect Dis. 2011 Jan;11(1):68.

[53] Van Cauteren D, Vaux S, de Valk H, Le Strat Y, Vaillant V, Lévy-Bruhl D. Burden of influenza, healthcare seeking behaviour and hygiene measures during the A(H1N1)2009 pandemic in France: a population based study. BMC Public Health. 2012 Jan;12:947.

[54] Biggerstaff M, Jhung MA, Reed C, Fry AM, Balluz L, Finelli L. Influenza-like illness, the time to seek healthcare, and influenza antiviral receipt during the 2010-11 influenza season – United States. J Infect Dis. 2014;.

[55] Jernigan DB, Lindstrom SL, Johnson JR, Miller JD, Hoelscher M, Humes R, et al. Detecting 2009 pandemic influenza A (H1N1) virus infection: availability of diagnostic testing led to rapid pandemic response. Clin Infect Dis. 2011 Jan;52 Suppl 1(Suppl 1):S36–43.

[56] Viboud C, Grais RF, Lafont BaP, Miller Ma, Simonsen L. Multinational impact of the 1968 Hong Kong influenza pandemic: evidence for a smoldering pandemic. J Infect Dis. 2005;192:233–248.

